# Is single nucleus ATAC-seq accessibility a qualitative or quantitative measurement?

**DOI:** 10.1101/2022.04.20.488960

**Authors:** Zhen Miao, Junhyong Kim

**Affiliations:** Graduate Group in Genomics and Computational Biology, Perelman School of Medicine, University of Pennsylvania, Philadelphia, PA, USA; Department of Biology, University of Pennsylvania, Philadelphia, PA, USA

## Abstract

Single nucleus ATAC-seq is a key assay for gene regulation analysis. Existing approaches to scoring feature matrices from sequencing reads are inconsistent with each other, creating differences in downstream analysis, and displaying artifacts. We show that even with sparse single cell data, quantitative counts are informative for estimating a cell’s regulatory state, which calls for consistent treatment. We propose Paired-Insertion-Counting (PIC) as a uniform method for snATAC-seq feature characterization.

## Main

Single nucleus ATAC-seq (snATAC-seq) assays open chromatin profiles of individual cells. However, unlike RNA-seq where the counts estimate numbers of molecules, there is not a common agreement on what biological state is being estimated from snATAC-seq data. Existing snATAC-seq analysis methods create chromosomal domain features either by arbitrarily dividing the entire genome into fixed-width segments (features usually referred to as bins), or estimating discrete domains by peak-calling from aggregated pseudo-bulk data (features usually referred to as peaks). Using bins as features has problems associated with arbitrarily fixing length scales and phase (i.e., starting positions of the bins) and the problem that many bins will contain no relevant information. Peaks subset functionally relevant genomic intervals, but there are technical challenges to resolve boundaries for heterotypic datasets and to identify functional elements for rare cells, and differences exist in numerical criterion for peak identification. After choosing bins or peaks, some methods assign the feature counts based on the number of fragments that overlap with a region (fragment-based counting; e.g., Signac^1^ and snapATAC^2^), while others assign counts based on the number of insertions within the region (insertion-based counting; e.g., 10X cellranger ATAC^3^ and ArchR^4^). After feature counting, most methods convert the counts into a binary state of “open” or “closed” (e.g., snapATAC^2^, SCALE^5^, scOPEN^6^, MASETRO^7^, and cisTopic^8^), while other retain quantitative count information, implying that single nucleus assays may contain quantitative information on nucleosome density or turnover (e.g., scABC^9^, chromVAR^10^, and ArchR^4^).

When considering counts, the configuration of fragment/insertion positions around the peak/bin interval can create different quantifications dependent on whether one uses fragments or insertions (Figure 1a-b). Histograms of counts for fragment-based or insertion-based counting applied to the same dataset (10X Genomics peripheral blood mononuclear cell dataset, PBMC-5k) show evident differences (**Figure 1c-f** and **Supplementary Table 1**). In particular, with insertion-based counting, there is an artifact of depleted odd numbers. In a standard ATAC-seq experiment, two Tn5 insertions in the appropriate directions are required to form one amplicon fragment, thus the unit of observation is pairs of insertions. Odd number of insertions only arise when rare fragments cross feature boundaries, artificially breaking up paired insertions of a fragment. Fragment-based counting also has problems because the entire interval of an amplicon from a pair of insertion is considered evidence of “openness”. However, longer the fragment, less likely the region away from the insertion sites is open. This is especially acute when there are long fragments with insertions completely outside the peak/bin of interest^11,12^ (cell 1 in **Figure 1a**). The two counting strategies can result in discrepancies in downstream analysis. As an example, we analyzed a P0 mouse kidney snATAC-seq dataset^13^ for Differentially Accessible Region (DAR) identification between two most abundant cell types with ArchR^4^ and Signac^1^ (**Methods**). We found up to 4.7% peaks are only significant with one counting strategy, but not the other (**Supplementary Figure 1a**).

**Figure 1.**
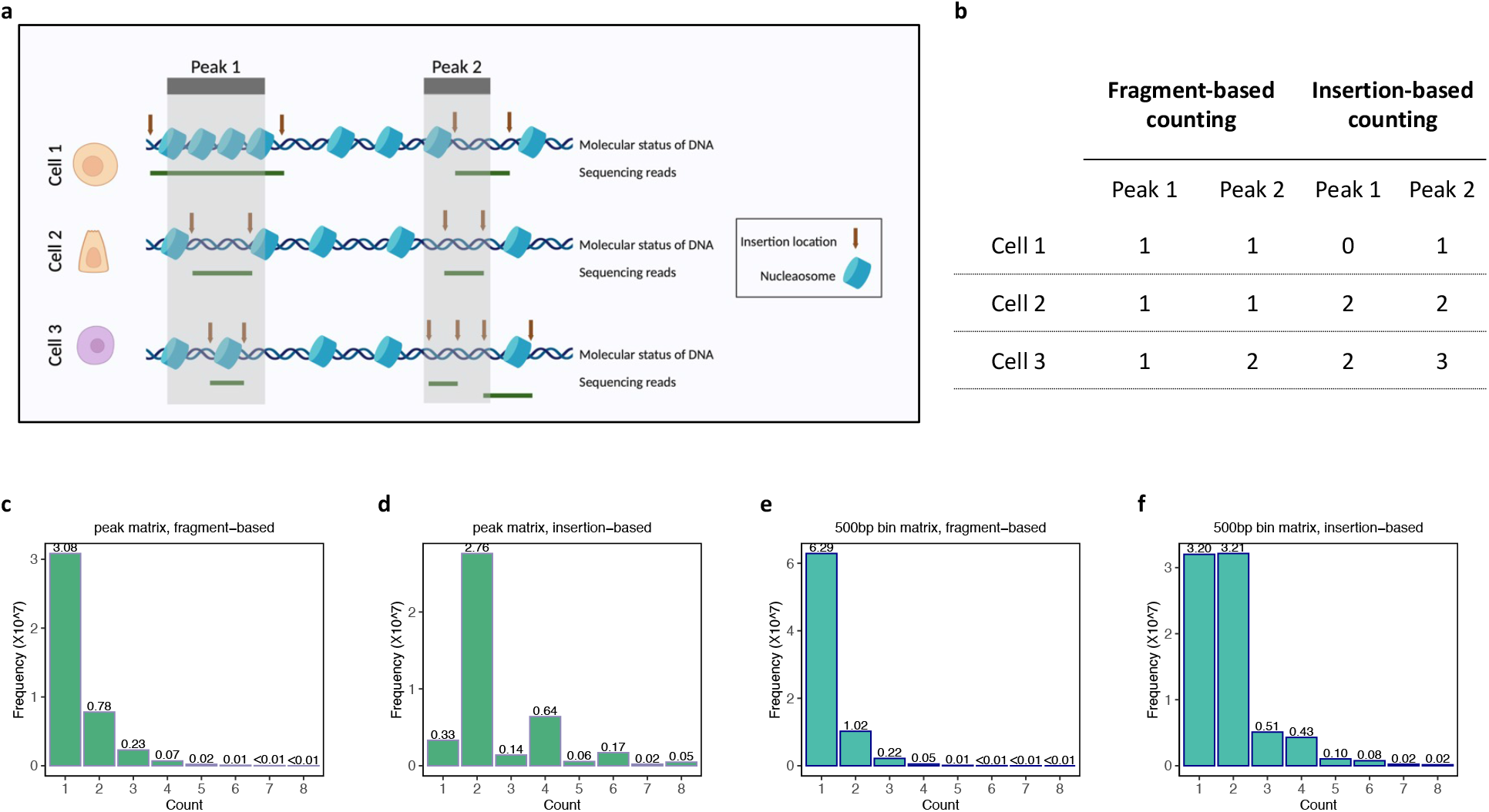
Two existing counting strategies for snATAC-seq data processing. (a-b) Schematic example of how the same open chromatin profiles can result in different counts with insertion-based or fragment-based counting strategies (c-f) Histogram of count frequencies with two counting strategies and with peaks or bins as features

If the counts are binarized, both insertion and fragment counting are consistent with each other, except for rare cases (e.g., cell 1 in **Figure 1a**). Thus, the vagaries of counting only matter if snATAC-seq contains quantitative information about the chromosome state. While variable nucleosome density and turnover dynamics imply that “openness” is a quantitative state^14^, it is not clear whether sparse data in single cells contain quantitative information. We asked whether more fragments in a peak for a single cell indicates higher probability that a randomly selected cell of the same type would be in open state. That is, we asked whether within-cell insertion density is predictive of between-cell sampling of open states. We first analyzed a human cell line snATAC-seq dataset^4^. The cell-by-peak matrix was constructed with insertion-based counting. We retained 166,142 peaks and 10,832 cells in ten cell types after stringent quality control (QC; see **Methods**). For each peak, we estimated the proportion of cells with the peak being accessible (hereafter we denote as open probability) in each of the ten cell types (**Methods**). With insertion-based counting approach, a count greater or equal to three indicates at least two fragments (four insertion events)— we call such cases “high density peaks”. We calculated the relative proportion of cells with high density peaks for each of the ten cell types (i.e., *P*(*y* ≥ 3|*y* > 0)) and then compared their rank order with the rank order of cell type open probability by Spearman rank correlation. Among the peaks we tested, the great majority (>94.6%) showed positive correlation and 9.4% showed significant correlations at significance level of 0.05 after FDR p-value correction (34.5% without FDR correction, **Figure 2a**). We also investigated the relationship between open probability and the relative proportion of cells with counts equal to two given counts being either one or two (i.e., *P*(*y* = 2|*y* = 1 *or* 2)) for the ten cell types. Consistent with our reasoning that the occurrence of one insertion mostly represents the boundary phasing artifact, we observed a symmetric distribution of Spearman correlation coefficients centered around 0 (**Figure 2b**), with only ∼0.08% peaks showing significant correlations at significance level of 0.05 after FDR p-value correction. Example peaks are shown in **Figure 2c-d** and **Supplementary Figure 1b-c**. We next examined the P0 mouse kidney snATAC-seq dataset^13^ we examined above. After QC, we retained 256,574 peaks and 9,286 cells in seven most abundant cell types in the dataset. With both insertion-based and fragment-based counting matrices, we conducted the same analysis as above, and the results were consistent with the human cell line data (**Supplementary Figure 2a-c**) where we found high-density peaks provided significant information on greater probability of open peaks in the corresponding cell type.

**Figure 2.**
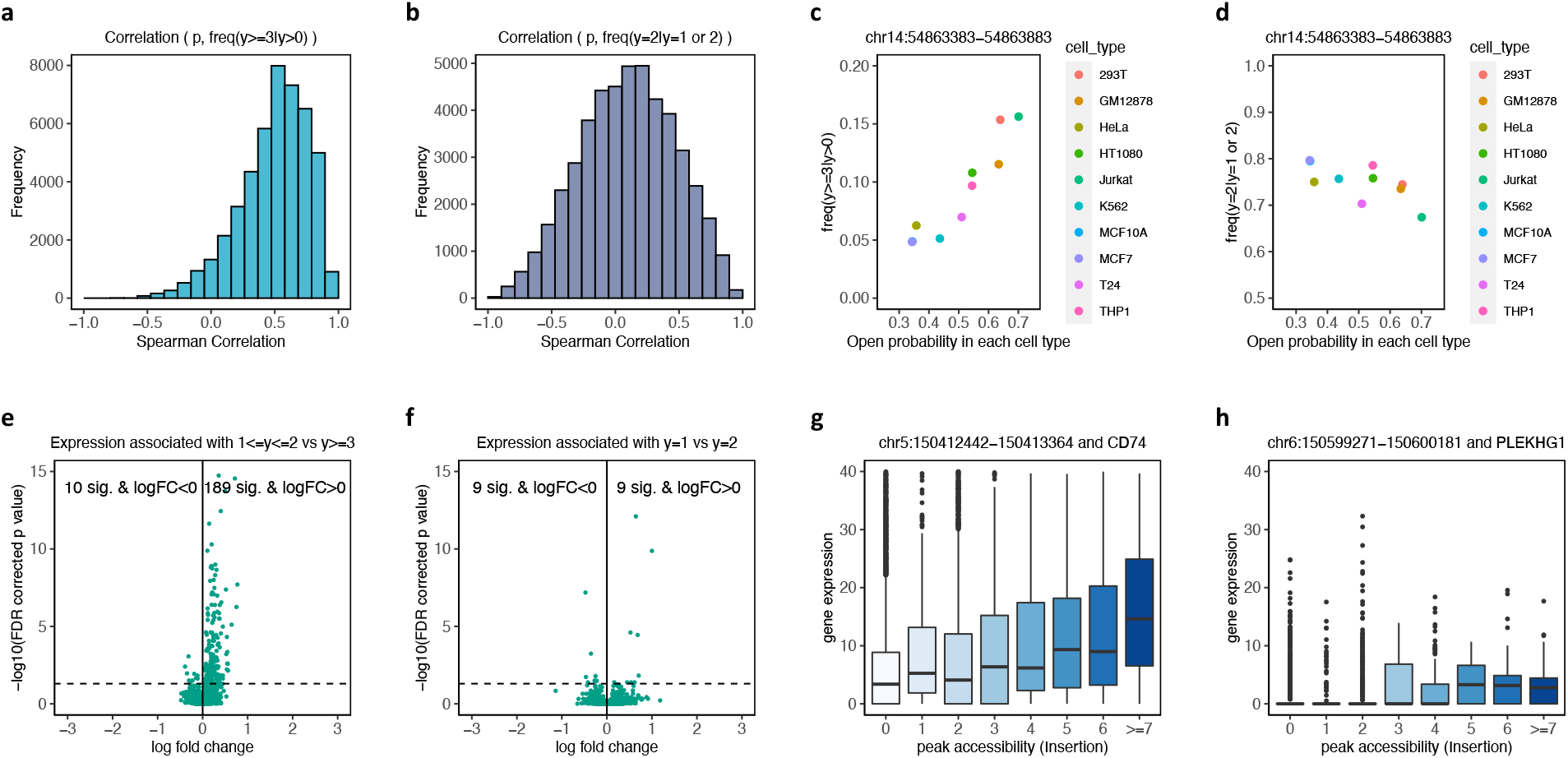
snATAC-seq data contain quantitative information of cellular states. (a) Histogram of Spearman correlation coefficients between open probability in each group and the relative frequency of counts greater than or equal to 3 in human cell line data (b) Histogram of Spearman correlation coefficients between open probability in each group and the relative frequency of counts equal to 2 given counts being either 1 or 2 in human cell line data (c-d) An example peak with different open probabilities across various cell types and the relative frequency of peaks with counts greater than or equal to 3 or the relative frequency of counts equal to 2 given counts were either 1 or 2 in human cell line data. Another example was displayed in **Supplementary Figure 1b-c** (e) Volcano plot showing the normalized gene expression levels between cells with TSS peak insertion counts equal to 1 or 2 and cells with TSS peak insertion counts greater than or equal to 3 in PBMC data (f) Volcano plot showing the normalized gene expression levels between cells with TSS peak insertion counts equal to 1 and cells with TSS peak insertion counts equal to 2 in PBMC data (g-h) Examples of peak-gene pairs where gene expression levels are related to the TSS peak insertion counts in PBMC data

To investigate the potential relationship between snATAC-seq count and gene expression, we analyzed a 10X genomics PBMC multiome dataset with RNA and ATAC measured on the same cells. We quantified the cell-by-peak matrix with insertion-based counting approach. Because the regulatory structure of chromatin domains around a given gene may be complex and largely unknown, we considered only peaks that are close (± 100 *bp*) to Transcript Start Site (TSS) to focus on the most proximal relationship. We also focused on peaks that had a broad range of one to four counts across cells, filtering out those with too small number of cells within appropriate range (**Methods**). This resulted in 3,387 peak-gene pairs across 11,234 cells. We compared the gene expression levels with associated TSS peak insertion count = 1 or 2 (single fragment) against those with count ≥ 3 (more than two fragments) using Wilcoxon rank sum test. We found 199 significant peak-gene pairs after FDR correction, 189 of which have positive log fold change (**Figure 2e**); 67.2% of peak-gene pairs showed higher non-zero expression proportion in the group of count ≥ 3. When we compared gene expression levels associated with TSS peak insertion count = 1 against those with count = 2, we found only 18 significant peak-gene pairs after FDR correction, nine of which have positive log fold change (**Figure 2f**). In addition, 52% peak-gene pairs showed higher non-zero expression proportion in the group of count = 2, suggesting no difference between the two groups. **Figure 2g-h** shows two examples of peak-gene pair where the distribution of RNA expression monotonically changes as a function of ATAC counts. We next analyzed a Bone Marrow Mononuclear Cells (BMMC) multiome dataset^15^ which again indicated that peak density was informative for expression levels (**Supplementary Figure 3a-d**).

In sum, greater counts of snATAC-seq insertions are correlated with greater probability of peak open state and higher expression of proximal genes, suggesting that even with single nuclear data, quantitative counting provides important functional information about the epigenomic state of the cell. We noted above that insertion-based counting creates occasional artifacts and ignores the fact that, while insertions themselves may be random, the sequence evidence is always in terms of pairs of insertions. Fragment-based counting has the problem that direct evidence of open state is only at the insertion site and the evidence for open state decays as a function of distance from the insertion site. Ideally, it might be appropriate to estimate the quantitative open state of an interval as a function of fragment lengths and local chromosome features. However, such a model will need to be data-driven given the irregularities of locus-specific chromosome dynamics. Here, we propose a simple consistent counting strategy we call Paired-Insertion-Counting (PIC, https://github.com/Zhen-Miao/PIC-snATAC). With PIC, for a given chromosome interval, if an ATAC-seq fragment’s pair of insertions are both within the interval, counted as one (pair); if only one insertion is within the interval also count one (pair).

PIC is consistent with the fact that all fragments have two insertions. It also prevents counting a fragment when its ends are both outside the peak/bin interval. It has the drawback that when one insertion is in the peak/bin and the other insertion is far from this insertion, evidence is weak that both insertions provide information on the current peak/bin. However, in most datasets, long fragments are rare and unlikely to greatly distort the data (**Supplement Figure 4**). We recommend treating snATAC-seq PIC count as a quantitative trait, wherever sensitivity is a critical factor.

In sum, snATAC-seq is increasingly an important tool for genomic analysis and despite sparse data at single cell resolution, we find evidence that it can be informative to consider “openness” as a quantitative trait. Existing approaches are inconsistent in how they quantify peak/bin openness and here we propose a new counting method that is consistent with the molecular basis of the assays.

## Methods

### Public Datasets

We downloaded the following snATAC-seq datasets from public repositories: mouse kidney data^13^ (GEO accession number GSE157079), human cell line data^4^ (GEO accession number GSE162690), and human BMMC data^15^ (GEO accession number GSE194122). We downloaded the 10X Genomics human PBMC data (including a snATAC-seq dataset and a sn-multiome dataset) from 10X Genomics website (https://www.10xgenomics.com/resources/datasets).

### Data QC and pre-processing

To remove artifacts due to data processing, we conducted QC filtering for the datasets. First, we removed peaks with very high counts (≥7 with fragment-based counting or ≥ 14 with insertion-based counting) across the entire dataset, which could be associated with repetitive or potentially uncharacterized blacklist regions^2^. We removed potential doublet cells by the number of regions with per-base coverage greater than 3 (Ref. ^16^). We also removed fragments with interval length smaller than 10 that are likely to be misalignment.

### Processing 10X Genomics PBMC snATAC-seq data (5k)

The 10X Genomics PBMC snATAC-seq data (ID: atac_pbmc_5k_nextgem) were used to compare the count distribution obtained from different counting methods. The peak ranges and insertion-based peak-by-cell count matrices were obtained from cellranger pipeline. The insertion-based bin-by-cell matrix was constructed by ArchR^4^. Bins that are accessible in fewer than ten cells were filtered. To obtain the fragment-based peak or bin count matrix, we used Signac^1^ pipeline.

### Adjusting Open Probability

We define “open probability” as the probability that a given genomic region is accessible for a randomly sampled cell of a given cell type. Note that this open probability does not measure the degree of openness but the probability of capturing a cell in an open state accessible to ATAC-seq assay. This probability will be governed by the temporal dynamics of nucleosome-dependent accessibility of that region for that cell type. Typical snATAC-seq data have missing data issue and are very sparse. In order to unbiasedly estimate the chromatin open probability in each cell type, we considered two sources of excessive zeros in the snATAC-seq data: biological inaccessibility and technical failure to capture open state in sequencing data. We developed the following model to estimate true open proportion.

Let 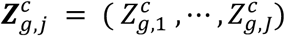 be a *J* × 1 binary vector denoting the open chromatin status of cell *c* that depends on group label *g* (e.g., cell type label). Each element in the vector, 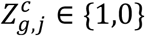 represents the accessibility of *j*^*th*^ genomic region (e.g., bin or peak), where the value 1 indicates open and 0 indicates close. We consider 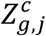 to be sampled from a Bernoulli distribution parameterized by *p*_*g,j*,_ the probability that a random cell of *g* type will be open for *j*^*th*^ region:

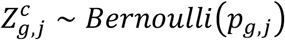

In practice, the true open chromatin status *Z* of cell *c* is unobserved. Instead, due to disparity of enzyme activity and sequencing depth across cells, an open state may not be observed in the data. We introduce 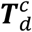 *J* × 1 binary vector representing the capture state of different enomic regions in cell *c*. This status depends on sequencing depth *d* for cell c. Additional experimental factors and the particular chromosomal region may also affect the status, which we ignore here. We also drop index *d*, since every cell is associated with particular sequencing depth. We assume:

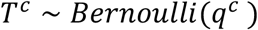

for some parameter vector *q*^*c*^ that is a function of the cell.

Let 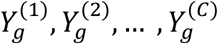 be a random vector representing observed data with *g* ∈ {1, 2, …, G} a priori assigned cell type labels. 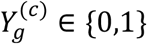 where 1 indicates open and 0 indicates close. Then 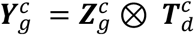 where ⊗ denote element-wise direct product (Hadamard Product).

For a given dataset ***y***, we set the loss function *log L*(***p, q***|*y*) as

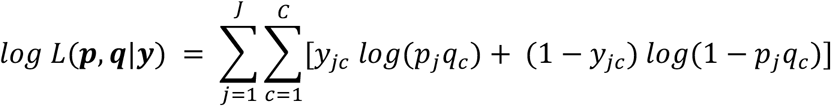

In order to compute both estimators for ***p*** and ***q***, we implemented a coordinate descent algorithm. This iteration stops until convergence:

1. Start with an initial estimate of ***p***^*(0)*^
2. For *t* = 1, 2, …
  a. Compute 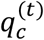 by:

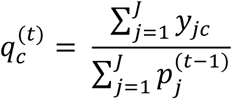
  b. Update 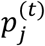 by moment estimator:

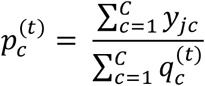

### Analysis of count frequency and open probability in human cell line data

The cell line data matrix was constructed by insertion-based counting method, and the maximum count was 4 in this matrix. The open probability for each cell type, *p*_*g*_ was estimated with the method described above. Since the count 2 and 1 mainly represent the boundary phasing issue, we estimated the probability of observing count greater or equal to 3 given observing a non-zero count, *P*_*g*_[*y* ≥ *3*|*y* > 0]

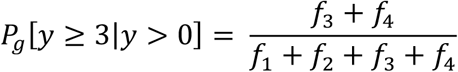

Since some peaks do not have counts that are greater than three, we only retained peaks with at least five count greater than 3, and 46,499 peaks were left. The Spearman correlation was computed between the open probability and frequency of counts greater than three. In addition, we also computed the probability of observing a count equal to 2 given the count being 1 or 2, *P*_*g*_[*y* = 2|*y* > 0]

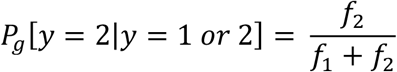

and its correlation with open probability.

### Analysis of differentially accessible regions (DAR) in P0 mouse kidney data

The peak information as well as cell type annotations were obtained from the original publication^13^. The peak-by-cell matrix was then constructed by both insertion-based and fragment-based approaches. The count correspondence is summarized in the **Supplementary Table 2**. We then picked the two most abundant cell types, nephron progenitor cells and stroma cells for the DAR analysis. Two DAR approaches, Signac^1^ and ArchR^4^, were used to identify DARs. Peaks with FDR-adjusted p value ≤ 0.05 were regarded as DARs.

### Analysis of count frequency and open probability in P0 mouse kidney data

We retained cell types with more than 600 cells to get accurate estimations of the parameters, which resulted in seven cell types. The open probability for each cell type, *p*_*g*_ was estimated with the method described above. Within a cell type, assuming there are *f*_1_ cells with count 1, *f*_2,_ cells with count 2 and so on, the probability of observing counts greater than or equal to 3 given observing a non-zero count is estimated by

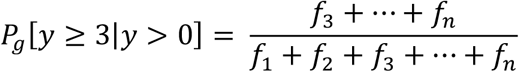

Spearman correlation was computed between the two quantities, and results were shown in **Supplementary Figure 2a-b**. We observed the same pattern with fragment-based counting when we compare the rank correlation between open probability and *P*_*g*_[*y* ≥ 2|*y* > 0]. (**Supplementary Figure 2c**).

### Analysis of gene expression and different counts for PBMC data

The 10X Genomics PBMC sn-multiome data (ID: pbmc_granulocyte_sorted_10k) were used to study the relationship between the number of insertions around TSS and its associated gene expression. We first retained peaks that overlap with ± 100 *bp* region around TSS and with at least five instances of counts greater than or equal to two. Then, we linked these peaks with their associated genes to form peak-gene pairs. The peak-gene pairs were then filtered by requiring the non-zero expression proportion with chromatin insertion counts greater than zero to be at least 10%. 3,387 such peak-gene pairs were kept for the downstream analysis.

For each peak-gene pair, we grouped the normalized gene expression levels by the insertion count in the TSS peak. Mean expression level and non-zero expression proportion were calculated for each group. Two-sided Wilcoxon Rank Sum test was then conducted between the two groups and log fold change was computed by comparing the mean expression differences.

### Analysis of gene expression and different counts for BMMC data

The BMMC dataset^15^ was collected across multiple institutes and multiple donors with batch effect. To prevent batch effect, we focused on one donor sample that was collected at one institute (donor #2 collected from institute #1). There are 6,740 cells across multiple cell types. With the same filtration criteria as above, we retained 2,488 peak-gene pairs for our analysis. The same analyses were conducted as above and were shown in **Supplementary Figure 3a-c**.

## Supplementary Information

**Supplementary Figure 1.**
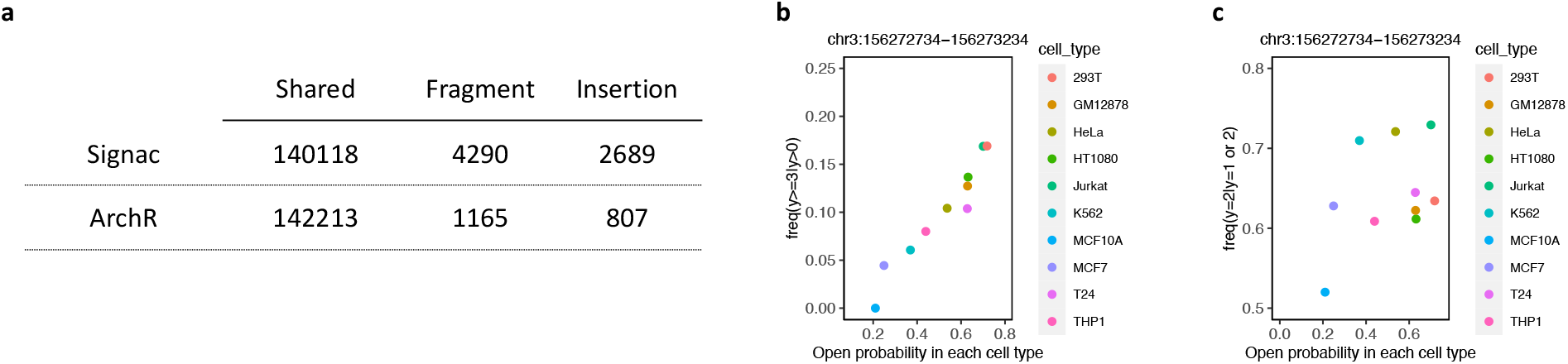
(a) Number of significant Differentially Accessible Regions between the two most abundant cell types, nephron progenitor cells and stroma cells with two different counting approaches and two different pipelines (b-c) An example of a peak with different open probabilities across various cell types and the relative frequency of peaks with counts greater than or equal to 3 or the relative frequency of counts equal to 2 given counts were either 1 or 2. Another example was displayed in **Figure 2c-d**

**Supplementary Figure 2.**
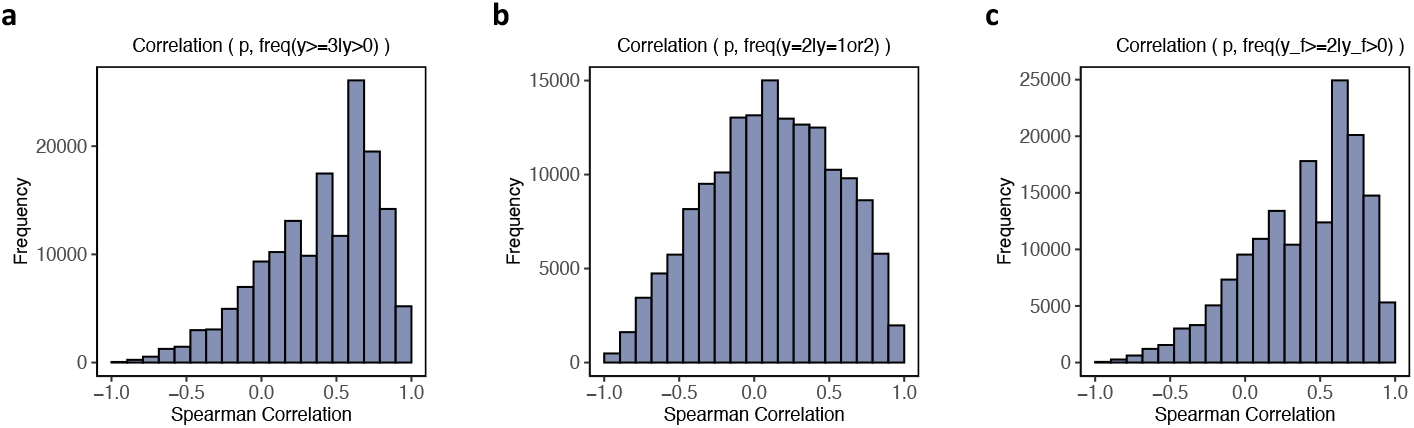
(a) Histogram of Spearman correlation coefficients between open probability in each group and the relative frequency of counts greater than or equal to 3 in P0 mouse kidney data (b) Histogram of Spearman correlation coefficients between open probability in each group and the relative frequency of counts equal to 2 given counts being either 1 or 2 in P0 mouse kidney data (c) Histogram of Spearman correlation coefficients between open probability in each group and the relative frequency of counts greater than or equal to 2 with fragment-based counting in P0 mouse kidney data

**Supplementary Figure 3.**
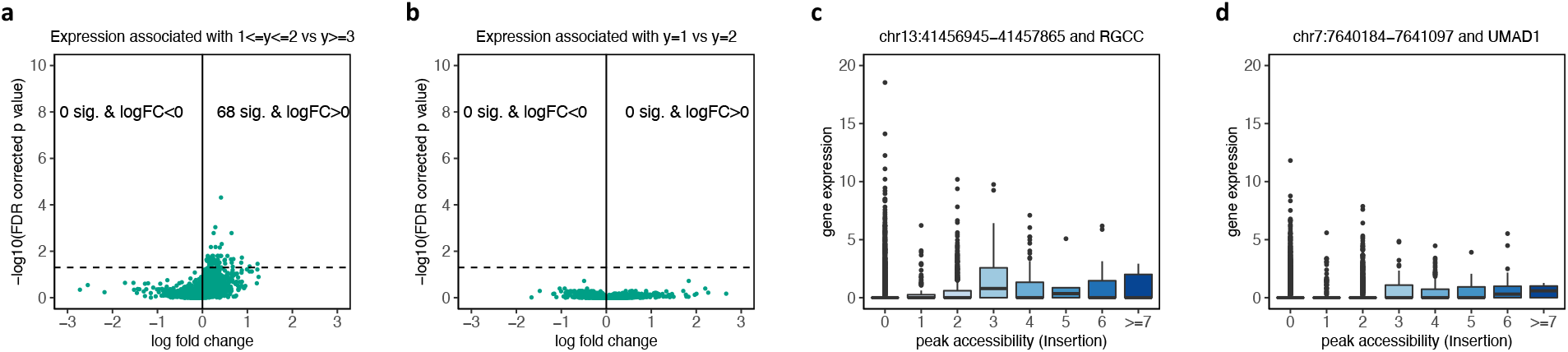
(a) Volcano plot showing the normalized gene expression levels between cells with TSS peak insertion counts equal to 1 or 2 and cells with TSS peak insertion counts greater than or equal to 3 in BMMC data (b) Volcano plot showing the normalized gene expression levels between cells with TSS peak insertion counts equal to 1 and cells with TSS peak insertion counts equal to 2 in BMMC data (c-d) Examples of peak-gene pairs where gene expression levels are related to the TSS peak insertion counts in BMMC data

**Supplementary Figure 4.**
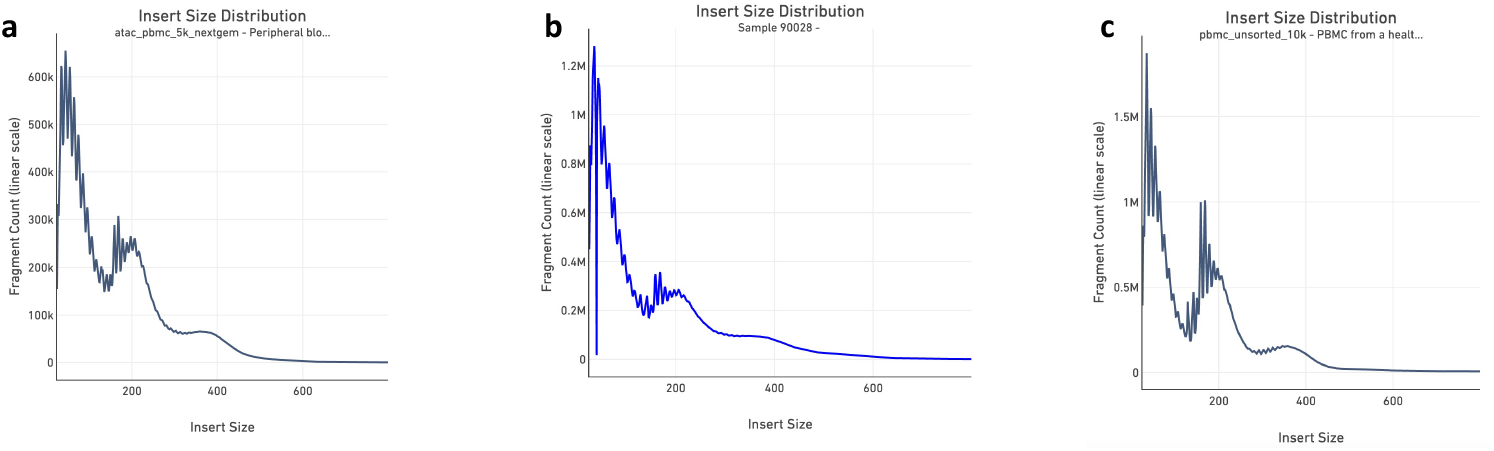
(a) Tn5 Insert size distribution in 10X Genmoics PBMC-5k snATAC-seq dataset (b) Tn5 Insert size distribution in P0 mouse kidney snATAC-seq dataset (c) Tn5 Insert size distribution in 10X Genmoics PBMC-10k snMultiome dataset

**Supplementary Table 1: Frequency of counts with different counting strategies (PBMC-5k data)**

**Supplementary Table 2: Correspondence between different counting strategies (kidney P0 data)**

## Acknowledgment

This work was supported by NIDDK grant 5UC2DK126024-02 as part of the ReBuilding a Kidney (RBK) consortium.

## Code Availability

All codes used in this project including PIC algorithm are in the GitHub repository: https://github.com/Zhen-Miao/PIC-snATAC

## Competing interests

The authors declare no competing interests.

## Notes

### Competing Interest Statement

The authors have declared no competing interest.

